# Systemic metabolic correlates of environmental sensitivity in group-housed mice

**DOI:** 10.64898/2026.05.18.725769

**Authors:** Natsumi Joko, Kentaro Abe

## Abstract

Environmental changes significantly impact the social behaviors of animals, yet individuals exhibit substantial variability in their responsiveness, known as environmental sensitivity. Understanding the biological basis of this individual variability is critical for elucidating vulnerability to stress-related and psychiatric disorders. To investigate the organism-wide physiological states linked to environmental sensitivity, we combined continuous, non-invasive RFID-based behavioral tracking with untargeted plasma metabolomics in group-housed mice undergoing spatial and social environmental restructuring. Following the environmental alteration, we observed heterogeneous behavioral shifts across individuals, enabling their operational classification into high-responsiveness mice (HRM) and low-responsiveness mice (LRM). Untargeted metabolomic profiling revealed distinct systemic metabolic signatures associated with these behavioral phenotypes. Specifically, HRM exhibited elevated levels of circulating essential and non-essential amino acids, as well as metabolites linked to one-carbon and energy metabolism. Exploratory co-variation analysis further identified plasma metabolic modules associated with individual behavioral metrics. These findings suggest that individual differences in behavioral adaptation are not solely neural phenomena but are coupled with coordinated, organism-wide metabolic adjustments. This study provides a framework for identifying candidate peripheral metabolic correlates of behavioral responsiveness to environmental and social change.

## Introduction

Behavior in social animals is strongly shaped by both physical and social environmental contexts (Sih *et al*, 2004; Coppens *et al*, 2010; Niemelä & Dingemanse, 2014). For instance, changes in habitat structure or spatial constraints can markedly impact social structures and territorial boundaries in animal populations (Blumstein *et al*, 2023). Human studies have similarly demonstrated that environmental architecture influences social connectivity and group dynamics (Sallis *et al*, 2006; Brown *et al*, 2014; Pinter-Wollman *et al*, 2018; Khogali *et al*, 2025). Importantly, individuals exhibit substantial variability in their responsiveness to such environmental changes (Belsky & Pluess, 2009; Wolf & Weissing, 2012). Maladaptation to environmental transitions is increasingly recognized as a contributing factor to the development of metabolic and psychiatric disorders (McEwen, 2007; Pyykkönen *et al*, 2010). For example, individuals who struggle to adjust to major life transitions, such as residential relocation or occupational change, are frequently diagnosed with adjustment disorder (Casey & Bailey, 2011; Luhmann *et al*, 2012). Likewise, patients with major depressive disorder (MDD) often exhibit reduced stress tolerance and decreased flexibility in response to environmental demands (Kendler *et al*, 1999; Casey & Bailey, 2011).

Investigating individual differences in behavioral adaptation using animal models presents methodological challenges. In particular, experimenter-induced disturbances can strongly influence behavior, especially under artificial or stressful testing conditions (Sorge *et al*, 2014; Segelcke *et al*, 2023). Conventional behavioral assays, which typically measure behavior at discrete time points in separate behavioral platforms, may introduce additional stressors and fail to capture the continuous and dynamic nature of social behavior (Datta *et al*, 2019). Recent technical developments have enabled continuous recording of group-housed animals using image-based tracking systems (Shemesh *et al*, 2013; Mathis *et al*, 2018; de Chaumont *et al*, 2019; Forkosh *et al*, 2019; Lauer *et al*, 2022; Fujibayashi & Abe, 2024). However, maintaining high tracking accuracy throughout experimental periods remains challenging, particularly when animals huddle together, move inside occluded areas such as shelters or obstacles, experience low-light conditions during the dark phases, or encounter environmental conditions that differ from those used for model training (Lauer *et al*, 2022). Moreover, in longitudinal studies assessing changes in behavioral patterns over extended periods, ID switching between animals can severely compromise the results (Peleh *et al*, 2019; Chen *et al*, 2023). Therefore, methodologies that ensure reliable individual identification are essential. These limitations make it difficult to assess how individuals adapt to environmental changes over time within a naturalistic social context.

Understanding the mechanisms underlying inter-individual variability is essential for elucidating vulnerability to environmental change across populations. Environmental sensitivity is unlikely to be solely a neural phenomenon but may instead involve coordinated physiological responses at the whole-organism level. Plasma metabolomics enables comprehensive profiling of circulating metabolites and provides a molecular interface linking behavioral phenotypes to peripheral physiological states (Nicholson *et al*, 2012; Patti *et al*, 2012). Given its low invasiveness, particularly in human studies, plasma metabolomic profiling has emerged as a promising approach for identifying molecular signatures associated with specific behavioral traits and neuropsychiatric vulnerability (Kaddurah-Daouk & Krishnan, 2009; Yang *et al*, 2020; Du *et al*, 2021; Yin *et al*, 2024; Cavaleri *et al*, 2026).

To obtain mechanistic insights into behavioral adaptation induced by environmental restructuring, we analyzed the behavioral dynamics of group-housed mice in response to alterations in cage geometry. We employed continuous RFID-based tracking to monitor the spatial positions and interactions of individual mice throughout the experimental period (Weissbrod *et al*, 2013; Klein *et al*, 2022; Lipp *et al*, 2024; Ochi *et al*, 2026). After establishing baseline social organization and positional preferences, the cage structure was modified to introduce a change in the social and physical environment. We found that these environmental changes induced pronounced behavioral reorganization across individuals. Notably, the magnitude of behavioral change varied among animals. Furthermore, these behavioral adaptations were accompanied by distinct alterations in plasma metabolic profiles, suggesting that peripheral metabolic states may reflect individual differences in responsiveness to environmental and social changes.

## Results

### Experimental environment for assessing behavioral change in group-housed mice

To examine social behavior at the group level, we housed four mice per cage in three independent groups and continuously tracked the location of each animal using a RFID-based floor antenna system (Ochi *et al*, 2026). Each mouse was implanted with a passive RFID tag in its abdomen prior to the experiments, enabling continuous localization with a temporal resolution of 0.1 s and a spatial resolution of 5 cm × 5 cm (a 6 × 8 grid array covering a 30 × 40 cm area) (Fig. 1A). Continuous tracking enabled simultaneous quantification of each mouse’s trajectory, revealing temporal dynamics in behavioral activity and spatial occupancy. (Fig. 1B–D). Social interactions could also be assessed from the tracking data of each mouse. For example, we quantified “invade-evade” events, defined as interactions in which one mouse (the invader) entered the adjacent grid cell of another mouse (Fig. 1E, F), causing the displacement of the latter mouse (Fig. 1G and H) (the evade response). These behavioral interactions have been used as a proxy for social dominance in previous studies (So *et al*, 2015; Battivelli *et al*, 2024; Zheng *et al*, 2025). Therefore, based on the frequency and directionality of invade-evade events, we estimated the social rank of each individual using the Glicko-2 rating algorithm, which exhibited dynamic changes throughout the experimental period (Fig 1I) (Glickman, 2012). Social hierarchy tended to be highly variable during the first several hours following group formation but gradually stabilized within approximately one day (Fig. 1I).

**Fig. 1.**
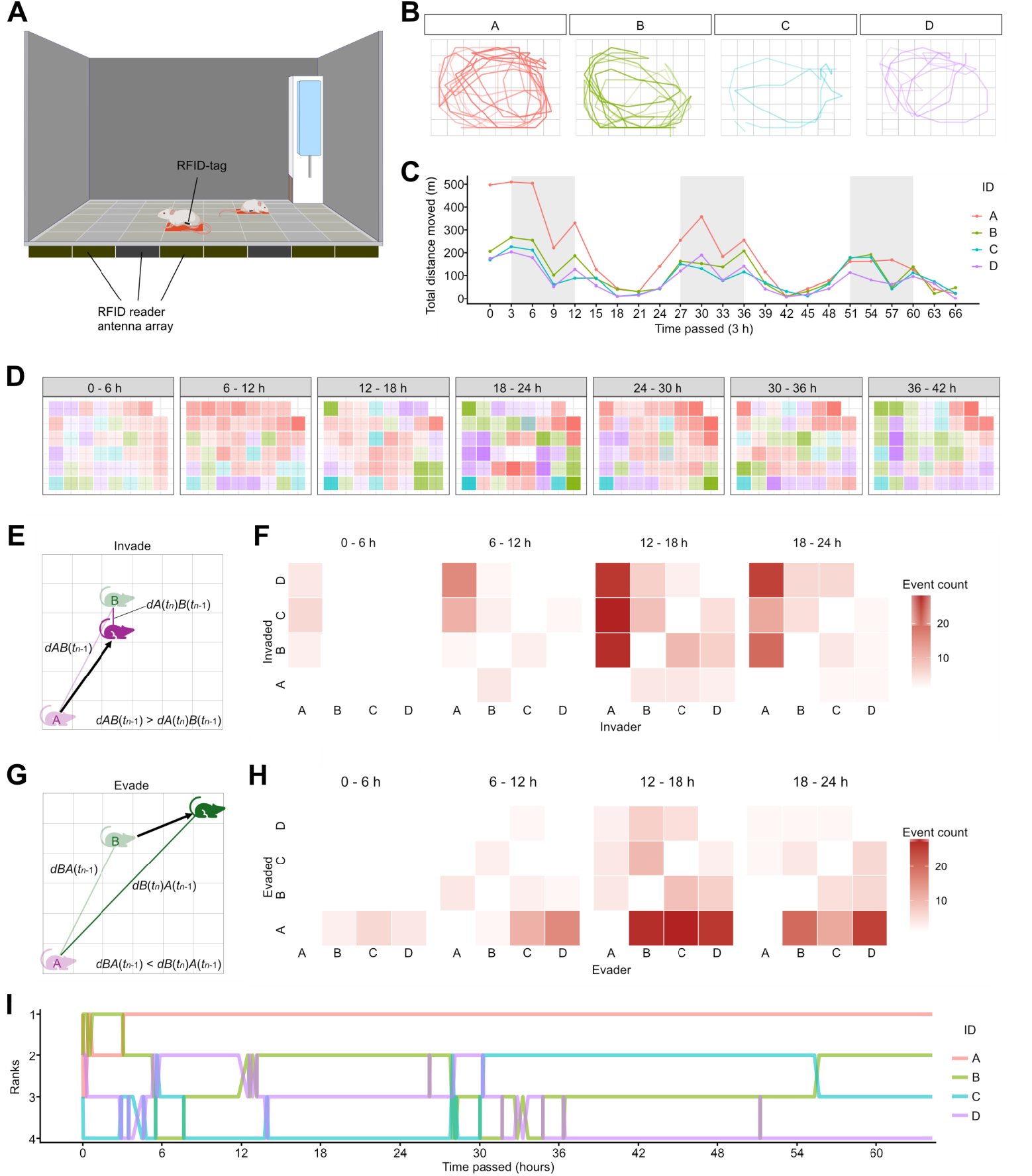
Longitudinal analysis of social behaviors in group. **(A)** Schematic of RFID-based mouse position tracking. Illustration was generated using BioRender. **(B)** Ten-minute trajectories of each mouse, with increasing opacity indicating the passage of time. **(C)** Total distance moved in 3-h bins. Shaded regions indicate dark phases; Colors indicate each animal. **(D)** Spatial occupancy in 6-h time bins. Color denotes the animal with the highest detection count in each spatial cell, and opacity indicates its proportional occupancy. **(E)** Schematic explaining the definition of Invade event. Invade is defined as the focal animal moving toward the target’s previous position. **(F)** Heatmap showing the number of directed invade events extracted from invade–evade interactions, from the focal (row) to the target (column) animal within each time bin. **(G)** Schematic explaining the definition of Evade event. Evade is defined as the focal animal moving away from the target’s previous position. **(H)** Heatmap showing the number of directed evade events extracted from invade–evade interactions, from the focal (row) to the target (column) animal during the corresponding time bin. **(I)** Time course of individual rankings computed from Glicko-2 ratings updated by invade–evade events.

### Environmental change induces heterogeneous social behavioral changes across individuals

Following the initial observation period, we introduced partitions to divide the cage into two compartments connected by corridors, thereby imposing an environmental change in spatial geometry (Fig. 2A). In parallel, one mouse was removed from each group to induce a change in social composition. These manipulations were introduced simultaneously to mimic a sudden environmental restructuring that alters both spatial organization and social composition. This manipulation produced profound changes in behavioral adaptations in the remaining mice. Some mice exhibited pronounced changes in spatial occupancy patterns and total movement distance, whereas others showed relatively little change (Fig. 2B). In addition, social hierarchy, quantified using Glicko-2 ratings derived from invade-evade events, exhibited dynamic reorganization following these manipulations, which stabilized within several hours (Fig. 2C and D). The behavioral changes observed across individuals were highly heterogeneous (Fig. 2E), suggesting differences in environmental sensitivity. We further analyzed the behavioral alterations of each mouse using 23 behavioral metrics extracted from the automated tracking data (Table S1) and calculated a change score for each individual by normalizing these metrics relative to baseline (Fig. 2F and G). This analysis identified mice exhibiting the largest and smallest behavioral shifts within the groups in which a significant difference in overall behavior was observed (ANOVA, F_(1, 22)_ = 5.55, group effect, *p* = 0.020). For subsequent analyses, mice exhibiting the largest behavioral shifts were operationally defined as high-responsiveness mice (HRM), whereas those showing minimal changes were categorized as low-responsiveness mice (LRM). Summing the squared Z-scores across all behavioral metrics revealed separation between the HRM and LRM groups (Fig. 2F). Consistently, principal component analysis (PCA) of these behavioral metrics revealed a separation between HRM and LRM (Fig. 2H). Further analysis of individual behavioral metrics indicated that HRM showed increased values in metrics associated with active behavior, whereas LRM exhibited relatively higher values in passive behavioral metrics (Fig. 2I). Collectively, these results demonstrate that behavioral dynamics changed following the environmental manipulation, with heterogeneous effects across individuals.

**Fig. 2.**
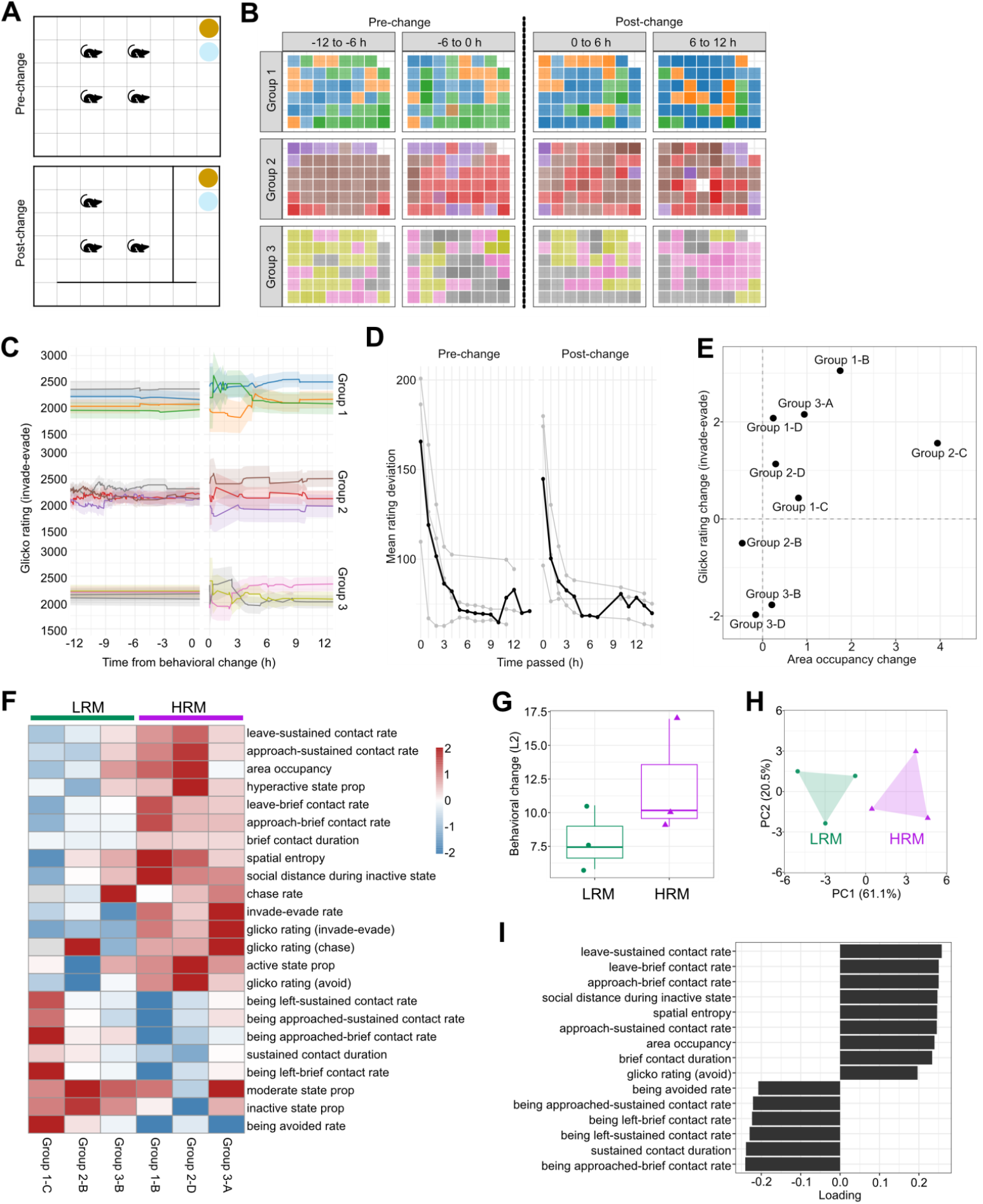
Behavioral changes followed by environmental change. **(A)** Schematical top-down view of the cage before and after the environmental change. Circles indicate zones for accessing food (brown) and water (light blue). **(B)** Spatial occupancy. Grid cells are colored by the extent of occupancy in 6-h bins before and after the environmental change. Color denotes the animal with the highest detection count in each spatial cell, and opacity indicates its proportional occupancy. **(C)** Time-course of Glicko-2 rating 12 h before and after environmental change. Colors denote each mouse. **(D)** Mean trajectories of individual Glicko-2 rating deviation (RD) for the three groups. The average across all three groups is indicated by a bold line. **(E)** Scatter plot showing the relationship between the average change in spatial occupancy and the average change in Glicko-2 rating (invade-evade) for each individual across groups. Dots represent individual values. **(F)** Heatmap showing z-scored changes across 23 behavioral metrics for each individual, grouped into HRM and LRM (*n* = 3 for each). **(G)** Dot and box plots show the L2 distances calculated from z-scored changes in all behavioral metrics. **(H)** Principal component analysis (PCA) of behavioral metrics across individuals. **(I)** Bar plot showing the contributions of individual behavioral metrics to PC1 (PC1 loadings).

### Plasma metabolomic profiles differ between high- and low-responsiveness mice

Given the pronounced inter-individual differences in behavioral responsiveness to environmental restructuring, we next examined whether these behavioral phenotypes were associated with systemic metabolic differences. To investigate this, we performed untargeted plasma metabolomic profiling following environmental manipulation. One day after the environmental change, mice were sacrificed and blood plasma was collected from six animals: three exhibiting pronounced behavioral changes (HRM) and three showing relatively minor behavioral changes (LRM). Untargeted metabolite profiling using capillary electrophoresis-mass spectrometry (CE-TOFMS), conducted in both cation and anion modes, detected a total of 271 metabolites across plasma samples (Fig. 3A and B). PCA demonstrated partial separation between HRM and LRM metabolic profiles, suggesting behavioral response-dependent metabolic remodeling (Fig. 3C). To identify coordinated metabolic changes, we applied weighted gene co-expression network analysis (WGCNA) to the plasma metabolomic dataset (Langfelder & Horvath, 2008). This approach clustered metabolites into six co-varying modules, each represented by a module eigengene (ME) (Fig. 3D–F). Comparative analysis revealed that three modules, ME-brown, ME-green and ME-magenta showed increasing trends in HRM relative to LRM, whereas ME-red showed decreasing trends (Fig. 3G). Pathway enrichment analysis of metabolites within the ME-green revealed a strong enrichment for central carbon and amino acid metabolism, including alanine, aspartate and glutamate metabolism, arginine biosynthesis, and the citrate cycle (TCA cycle). The ME-turquoise module was enriched for one-carbon and amino acid metabolic pathways, whereas the ME-magenta module was primarily associated with glycerophospholipid metabolism (Fig. 3H). Consistent with these findings, plasma concentrations of both essential and non-essential amino acids, as well as metabolites associated with one-carbon metabolism and energy synthesis, were significantly increased in the HRM group compared with the LRM group (Fig. 3I). These analyses highlighted a subset of metabolites within behaviorally associated modules that showed consistent relationships with specific behavioral responses to environmental restructuring.

**Fig. 3.**
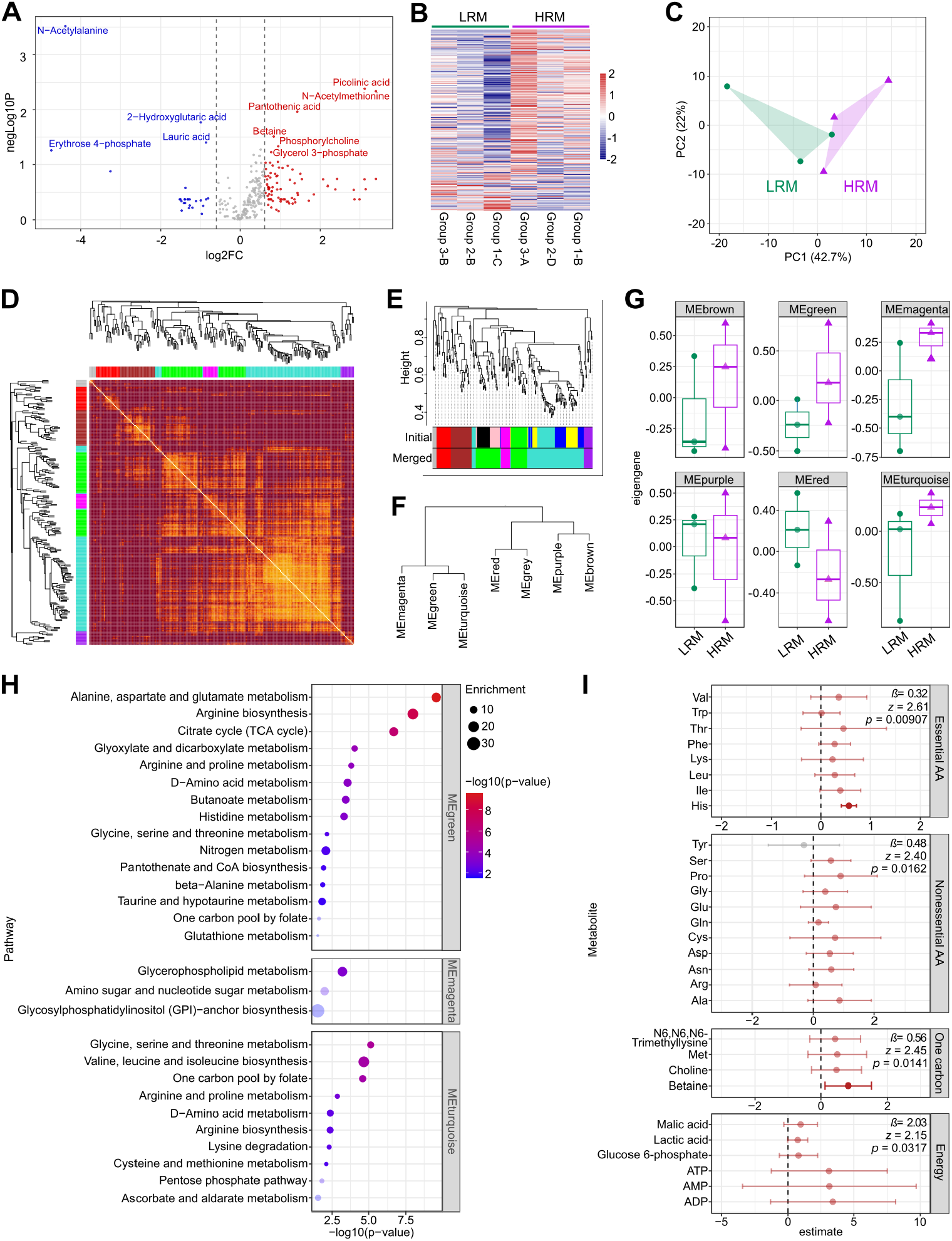
Metabolic difference between mice group of different sensitivity. **(A)** Volcano plot showing differential metabolite abundance between the HRM and LRM groups. The log_2_ fold change is plotted against the −log_10_(*p*-value). Dashed vertical lines indicate the fold-change threshold (|log_2_FC| = 0.6). Metabolites exceeding both fold-change and *p*-value thresholds (*p* < 0.1) are labeled. **(B)** Heatmap of metabolite abundance across samples. Metabolites are ordered by decreasing log2 fold change between the HRM and LRM groups without clustering. **(C)** Principal component analysis of log-transformed metabolite profiles. **(D)** Topological overlap matrix (TOM) heatmap from WGCNA network analysis. Metabolites are clustered based on network connectivity, revealing modules of highly co-regulated metabolites. (**E, F**) Hierarchical clustering dendrogram of metabolites showing module assignments before and after module merging (E) and the relationships among modules (F). **(G)** Distribution of module eigengenes. Each panel shows the eigengene of a WGCNA module. Points represent individual samples, and boxplots summarize the distributions within HRM and LRM groups. **(H)** Pathway enrichment analysis of metabolites within WGCNA modules. Point size represents enrichment ratio (observed/expected), and color indicates statistical significance. **(I)** Mixed effects models were fitted for each metabolite category with group as a fixed effect and metabolite and sample as random effects. Points indicate estimated group effects (*β*), and horizontal bars represent 95% confidence intervals. Red points indicate effects whose confidence intervals do not cross zero.

### Candidate metabolic modules covary with behavioral responsiveness

Because these metabolomic modules differed between high- and low-responsiveness individuals, we next examined whether they were quantitatively associated with behavioral variation across individuals. Specifically, we investigated whether metabolic modules were associated with behavioral features that contributed most strongly to the separation between HRM and LRM. To this end, we performed correlation analyses between module eigengenes and the first principal component (PC1) derived from PCA of 23 behavioral metrics. Linear regression analysis revealed a positive association between behavioral PC1, which captured the major axis of behavioral variation distinguishing HRM from LRM, and the ME-magenta module (*β* = 0.096, *p* = 0.014), with additional positive but non-significant associations observed for ME-green (*β* = 0.055, *p* = 0.37) and ME-turquoise (*β* = 0.078, *p* = 0.076) (Fig. 4A). When individual behavioral metrics were analyzed, we observed a general pattern of opposing correlations between modules enriched in HRM (ME-green, ME-magenta, and ME-turquoise) and those reduced in HRM (ME-red), suggesting coordinated but directionally distinct metabolic changes associated with behavioral variation (Fig. 4B). To further identify behaviorally relevant metabolites, we defined hub metabolites as those exhibiting high module membership (|kME| > 0.9) together with strong correlations with specific behavioral metrics (|*β*| > 0.8) (Fig. 4C). This analysis identified several metabolites that were strongly correlated with discrete behavioral indices (Fig. 4D). For instance, valine exhibited positive correlations with behaviors such as “approach-brief contact rate”, “leave-brief contact rate” and “social distance during inactive state”, while showing a negative correlation with “being approached-brief contact rate” (Table S1). Collectively, these findings indicate that specific plasma metabolic modules covary with behavioral phenotypes that emerge following environmental adaptation. These results suggest that systemic metabolic remodeling accompanies individual differences in behavioral responsiveness to environmental change.

**Fig. 4.**
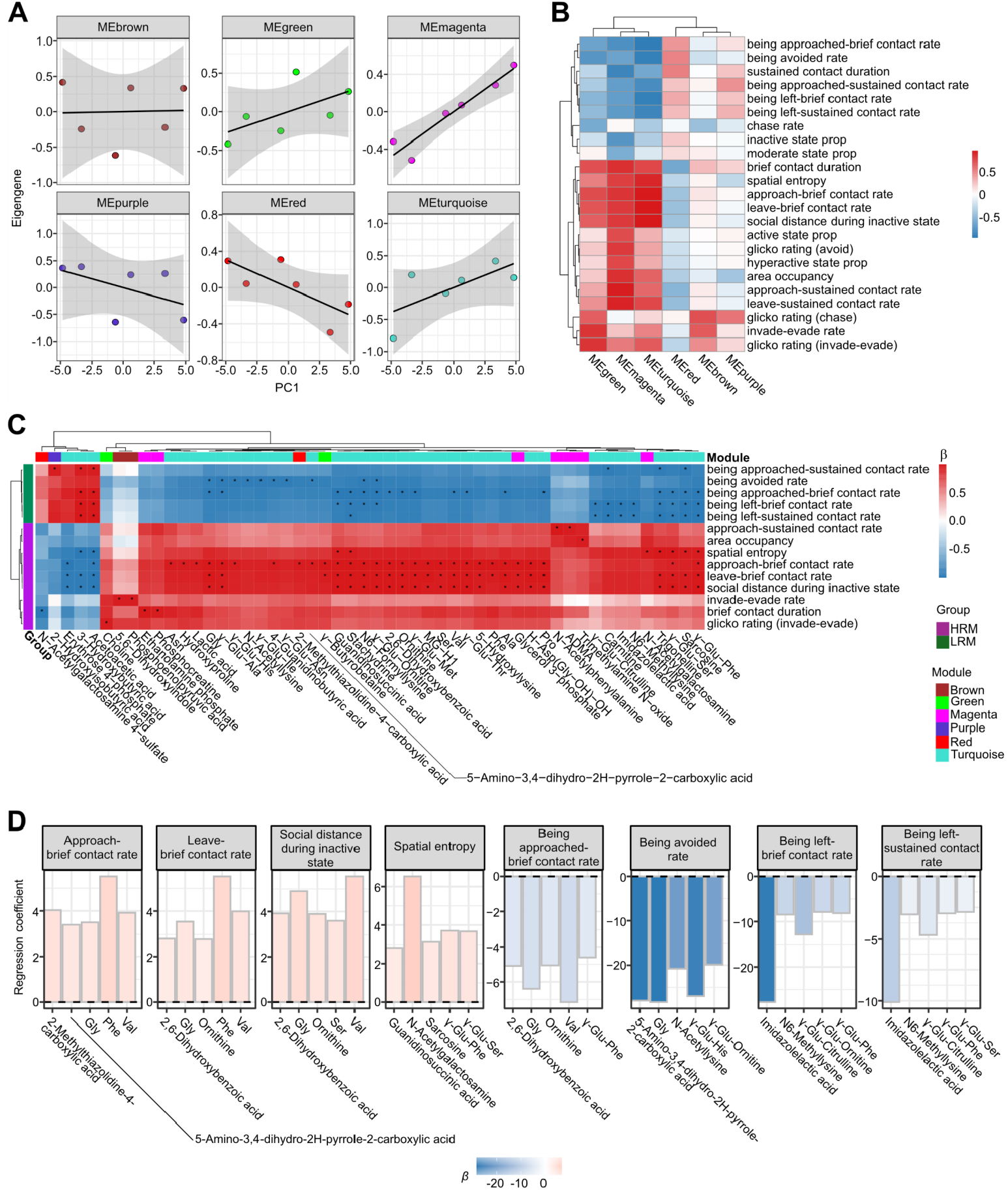
Relation between metabolites and behaviors. **(A)** Relationship between primary principal component (PC1) of 23 behavioral metrics and metabolic modules. Each point represents an individual. Lines indicate fitted linear regression with 95% CI (gray). **(B)** Heatmap summarizing associations between metabolite modules and 23 behavioral metrics. Colors represent the median standardized regression coefficient (*β*) across metabolites within each module. **(C)** Heatmap displaying standardized regression coefficients for associations between 14 behavioral features and each metabolite. Only subset of metabolite–behavior pairs are shown, filtered based on significance after multiple-testing correction and additional module-based selection criteria. Specifically, metabolites with strong module membership (|kME| > 0.9) and large effect sizes (|*β*| > 0.8) are shown. Asterisks indicate FDR-significant associations (*p* < 0.05). The top row denotes metabolite module identity, shown in colors. The leftmost column indicates the behavioral features most strongly associated with the HRM or LRM group. **(D)** Top five metabolites for each behavioral metric. Bars represent regression coefficients (*β*) obtained from linear models relating behavioral metrics to group batch-adjusted metabolite expression residuals. Bars are ordered by effect size, and colors indicate the direction and magnitude of the association.

## DISCUSSION

This study demonstrates that environmental restructuring within mouse social groups elicits substantial behavioral reorganization, with pronounced inter-individual variability in the magnitude of these responses. Untargeted plasma metabolomic profiling revealed distinct post-manipulation metabolic signatures associated with these behavioral differences. Notably, coordinated shifts in specific metabolic modules were associated with defined behavioral metrics, indicating that systemic metabolic remodeling parallels individual differences in behavioral adaptation. Given the exploratory nature of the metabolomic cohort, we focused on effect-size patterns, pathway-level trends, and candidate metabolite–behavior associations rather than definitive biomarker discovery.

Using automated and non-invasive approaches to quantify individual behaviors within mouse social groups, we were able to monitor behavioral dynamics without direct experimenter intervention. By minimizing experimenter-induced influences, this approach is particularly advantageous for assessing behavior under stressful conditions, such as the environmental perturbations examined in this study. In addition, this methodology enables continuous, longitudinal tracking of social behavioral dynamics, which is difficult to achieve with conventional analyses performed at discrete time points. Our findings also relate to a growing body of literature describing stable individual differences in behavioral responsiveness to environmental challenges (Sih *et al*, 2004). Previous studies have shown that animals differ consistently in their coping styles and stress responsiveness, often characterized as proactive versus reactive behavioral strategies (Coppens *et al*, 2010; Solié *et al*, 2026). Such individual differences, sometimes referred to as behavioral syndromes or animal personality traits, influence how individuals respond to environmental perturbations and social challenges (Wolf & Weissing, 2012). The heterogeneous behavioral responses observed in our study may reflect similar underlying variability in environmental sensitivity.

Our metabolomic analyses suggest that these behavioral differences are accompanied by coordinated systemic metabolic changes. In this study, because plasma metabolomes were compared across individuals, we could not determine whether the observed metabolic differences arose as a consequence of behavioral changes or reflected pre-existing molecular differences before the environmental reconstruction.

Longitudinal within-individual metabolomic analyses will be required to address this limitation and clarify the causal relationship between molecular state and behavioral adaptation. Despite this limitation, we identified clear biological differences between mice that exhibited significant behavioral changes and those that did not. Notably, HRM showed elevated levels of both essential and non-essential amino acids, which may reflect increased metabolic demand associated with enhanced locomotor activity and possibly enhanced social engagement following environmental restructuring (Takeshita *et al*, 2012; Axsom *et al*, 2024). Amino acids are important substrates for energy metabolism and gluconeogenesis and may also be mobilized through glucocorticoid-associated protein catabolism (Torres *et al*, 2023). In addition, several amino acids serve as precursors for neurotransmitters, raising the possibility that systemic metabolic changes may contribute to neurochemical adaptations accompanying behavioral responses to environmental perturbations (Fernstrom, 2013). The concurrent enrichment of one-carbon and purine metabolic pathways further suggests activation of biosynthetic and regulatory metabolic programs (Ducker & Rabinowitz, 2017). Collectively, these findings support the view that strong behavioral responsiveness to environmental change is accompanied by coordinated systemic physiological adjustments, rather than neural processes alone (Gautam *et al*, 2015; Papageorgiou *et al*, 2023). Although the causal relationship between metabolic state and behavioral sensitivity remains to be determined, our results suggest that peripheral metabolic profiles may provide measurable correlates of individual differences in environmental adaptability.

The extensive metabolic remodeling in HRM suggests strong mobilization of physiological resources, potentially reflecting increased metabolic demand alongside stress-associated catabolic processes. While the pronounced behavioral plasticity observed in HRM may enable rapid adaptation to abrupt environmental change, such heightened environmental sensitivity may also impose substantial biological costs. In line with the concept of allostatic load (McEwen, 2007), prolonged or repeated activation of such responses may lead to cumulative physiological burden. We therefore propose that high environmental sensitivity may act as a double-edged sword: although it may initially support active coping, the accompanying metabolic expenditure could predispose individuals to maladaptive outcomes when environmental stress is persistent. This framework offers one possible interpretation for the increased vulnerability of highly sensitive individuals to stress-related metabolic and psychiatric disturbances, including adjustment disorders and major depressive disorder (Kendler *et al*, 1999; Pyykkönen *et al*, 2010). In this view, the distinct metabolic signatures observed in HRM may reflect not only adaptive physiological engagement but also the early emergence of systemic burden, raising the possibility that they could serve as candidate biomarkers of vulnerability (Yamamoto *et al*, 2026).

In summary, our findings suggest that individual differences in behavioral responsiveness to social and spatial environmental restructuring are accompanied by distinct peripheral metabolic profiles. The observed coupling between behavioral state and plasma metabolic composition supports the idea that individual variability in environmental responsiveness is reflected in organism-wide physiological processes. Although exploratory, these results provide a framework for investigating peripheral molecular correlates of individual differences in behavioral adaptation. Future research comparing murine plasma metabolic profiles with human samples under analogous experimental conditions may provide translational insights and help bridge the gap between mouse models and human applications. Because peripheral metabolites can be measured with relatively minimal invasiveness, this framework may guide future translational studies of the physiological correlates of environmental sensitivity.

### Limitations of this study

Several limitations of this study should be acknowledged. First, the sample size for untargeted plasma metabolomic analysis was relatively small. Therefore, the metabolomic findings should be interpreted as exploratory, and future studies with larger, independent cohorts are required to validate the identified systemic metabolic signatures. Second, because blood samples were collected only after environmental manipulation, we cannot determine the causal direction between metabolic profiles and behavioral changes. Longitudinal metabolic profiling within the same individuals will be necessary to ascertain whether specific baseline molecular profiles predict environmental sensitivity or emerge as physiological consequence of behavioral adaptation. Third, our environmental manipulation simultaneously altered both spatial geometry and social composition by introducing partitions and removing a cage-mate, respectively. While this dual-manipulation paradigm successfully mimicked complex real-world transitions and maximized phenotypic divergence, it precludes us from dissociating the specific metabolic contributions of physical versus social stressors.

Future experiments employing isolated manipulations will be needed to dissect the precise drivers of these organism-wide physiological adaptations. Finally, our analysis was conducted only in male subjects. Further experiments in female mice will be required to generalize our findings.

## Materials and Methods

### Animals and animal care

Mice (*Mus musculus*, Slc: ICR) were obtained from SLC Japan and maintained in our facility under standard conditions, including housing in clear plastic cages in a temperature- and humidity-controlled room with a 12-h light/dark cycle (light on at 8:00 a.m. and off at 8:00 p.m.) with *ad libitum* access to standard food and water. The care and use of animals in this study were reviewed and approved by the Institutional Animal Care and Use Committee of Tohoku University (2018LsA-012, 2018LsA-013, 2020LsA-003, 2020LsA-004, and 2024LsA-002). All experiments and maintenance were performed following relevant guidelines and regulations.

### Behavioral analysis

A total of 12 male mice were used for behavioral analysis (*n* = 4 per group). One week prior to behavioral testing, each mouse was anesthetized using balanced anesthesia (medetomidine 30 µg/mL, All Japan Pharma; midazolam 30 µg/mL, Astellas Pharma; butorphanol tartrate 500 mg/mL, Meiji Seika Pharma; NaCl 118 mM; 400 µL per mouse), and a passive RFID tag (7 mm × 1.25 mm, Phenovance) was surgically implanted subcutaneously in the abdominal region. Mice were then allowed to recover for one week in a group-housed condition. For behavioral tracking analysis, mice were group-housed in a cage (size, 30 × 40 cm; height, 30 cm) made from gray PVC (5 mm thickness) to match the size of RFID reader antenna board (eeeHive 2D, Phenovance) which was placed beneath the cage. Food and water were placed adjacent to this cage, each accessible to the mouse through a single grid. For environmental manipulation, one randomly selected individual was removed from each group, and additional partitions were introduced into the cage. Twenty-four hours after the environmental manipulation, whole blood samples were collected from the remaining three mice under anesthesia.

### Behavioral tracking

The location of the RFID tag implanted in each mouse was detected by a 6 × 8 grid array of RF detectors within the antenna board hardware, which recorded the ID of the antenna detecting the strongest signal at each time point. Data were collected at 10 Hz throughout the experimental period (> 4 days). The collected data were transferred offline and analyzed using a custom program written in R. To track each mouse, we first reduced the temporal resolution to 2 Hz. Then, we assigned each mouse’s position to *X*–*Y* coordinates. Missing data points were imputed using the coordinates from the most recent prior detection. Movement was defined as displacement beyond an adjacent grid cell, or as a net displacement of at least two grid cells over three time steps.

### Definition of behaviors

Activity was quantified as the proportion of time spent in each state. Contact was quantified by its duration, whereas other dyadic interactions were evaluated by their counts. For all dyadic interactions, durations and counts were normalized by observation time and the number of individuals to derive rates relative to behavioral opportunities. In addition, invade–evade, chase, and avoid events were treated as pairwise interactions and analyzed using a Glicko-2 rating framework to estimate individual dominance. For each interaction, the winner was defined as the invader in invade–evade events, the chaser in chase events, and the individual being avoided in avoid events, while the counterpart was assigned a loss. Ratings were initialized at 2,200 with a rating deviation (RD) of 300 and a volatility parameter of 0.06 and updated sequentially across events using the Glicko-2 algorithm, allowing individual dominance estimates to vary over time. Behavioral metrics were aggregated in 6-h time bins. Behavioral changes were calculated as deviations from the pre-change baseline according to the following equation:

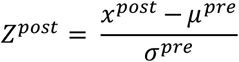

Changes in Glicko-2 rating were quantified as the difference between post- and pre-phase ratings (ΔRating). To account for uncertainty, a standardized change score (Δz) was calculated by dividing ΔRating by the pooled rating deviation across phases according to the following equation:

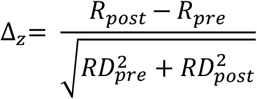

The definitions of the behavioral metrics are detailed in Table S1. Each behavioral metric was further revisualized by experimental batch to account for batch effects.

### Metabolome analysis

Each mouse was deeply anesthetized with 3.5 % isoflurane (Viatris Japan) inhalation. Whole-blood samples were collected by cardiac puncture using a syringe supplemented with EDTA-2Na, with the volume of blood collected adjusted to achieve a final EDTA concentration of 0.13%. The collected blood was immediately centrifuged at 1,200 ×g for 10 min, and the plasma was separated and stored at −80 °C until CE-MS analysis. CE-MS analysis was operated by HMT technology (Japan), using a CE-TOFMS system (Agilent Technologies). Both cation mode and anion modes were analyzed. Peak signals and m/z values were obtained using MasterHands software and aligned to registered metabolites.

Acquired metabolomic data were analyzed using R (v4.4.1). Relative abundance data were obtained from metabolite profiling and processed prior to downstream analyses. Missing values were replaced with half of the minimum non-zero value detected for each metabolite, followed by log_2_ transformation to stabilize variance. The following analyses were performed on metabolomic data: **Multivariate analysis**. Principal component analysis (PCA) was performed using centered and scaled data to assess global metabolic variation. Group separation was visualized using convex hulls. **Linear mixed-effects modeling**. To evaluate pathway-level changes, metabolites were grouped into predefined functional categories (e.g., essential and non-essential amino acids, one-carbon metabolism, and energy metabolism). Linear mixed-effects models were fitted using metabolite and sample as random effects according to following equation:

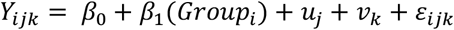

where *Y*_*ijk*_ denotes the metabolite abundance, *Group*_*i*_ represents the HRM or LRM phenotype, *β*_0_ is the intercept, *β*_1_ is the fixed group effect, *u*_*j*_ and *v*_*k*_ are random effects for the metabolite and sample, respectively, and *ε*_*ijk*_ is the residual error. Statistical significance was determined using Wald tests, and *p*-values were adjusted using the Benjamini–Hochberg method. **Weighted metabolite co-expression network analysis (WGCNA)**. Weighted correlation network analysis was performed using the WGCNA package (v1.73). Metabolites detected in fewer than 50% of the samples were excluded from this analysis. A signed adjacency matrix was constructed using a soft-thresholding power of *β* = 6, selected based on approximate scale-free topology criterion. A topological overlap matrix (TOM) was computed, and hierarchical clustering was performed using average linkage. Modules were identified using dynamic tree cutting (deepSplit = 2, minimum module size = 10), followed by module merging with a cut height of 0.3. Module eigengenes (MEs) were calculated as the first principal component of each module. **Pathway enrichment analysis:** This analysis was performed using the web-based platform MetaboAnalyst (v6.0; https://www.metaboanalyst.ca). Metabolites were mapped to HMDB identifiers and analyzed using over-representation analysis (ORA) with hypergeometric testing. *P*-values were adjusted using the false discovery rate (FDR), and pathways with an FDR < 0.1 were considered significant. **Integration with behavioral data**. Behavioral features, including principal component scores (PC1) and individual behavioral metrics, were summarized per individual and experimental condition. Residuals were calculated after adjusting for batch (date) effects. Associations between metabolite abundance and behavioral variables were evaluated using linear regression models according to following equation:

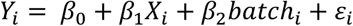

where *Y*_*i*_ represents the response variable (behavioral metrics, PC1, or module eigengenes), *X*_*i*_ represents the predictor variable (metabolite abundance or behavioral variables), *batch*_*i*_ is a covariate included to account for experimental batch (date) effects, and *ε*_*i*_ denotes the residual error. Standardized effect sizes (*β*) and corresponding *p*-values were obtained, and multiple testing correction was applied using the Benjamini– Hochberg method.

### Quantifications and statistical analysis

All statistical analyses were performed using R (4.4.1). Multiple-comparison correction was applied using the Benjamini–Hochberg method where appropriate. Statistical significance was defined as *p* < 0.05 or adjusted *p* < 0.05, as appropriate. Detailed information on the statistical analyses used for each experiment is provided in the corresponding Methods sections and figure legends

## Supporting information

Supplemental infomation

## Acknowledgments

We would like to thank Osumi N. and Ochi S. for sharing unpublished information regarding the RFID system with us. We thank Abe lab members for kind support during this study.

## Funding

This research was supported by JSPS/MEXT KAKENHI (JP25H010370, JP24H021460, JP24H012180 to K.A.), Tohoku University Research Program “Frontier Research in Duo” (Grant No. 2101 to K.A.), Tohoku University Neuro Global Program to NJ, Suzuken Memorial Foundation (K.A.), Takeda Science Foundation (K.A.), and Asahi Glass Foundation (K.A.).

## Author Contributions

Conceptualization: KA and NJ; Methodology: NJ; Investigation: NJ; Visualization: NJ; Data analysis: NJ; Funding acquisition: KA; Project administration: KA; Writing – original draft: KA and NJ; Writing – review & editing: KA and NJ.

## Competing interests

The authors declare no competing interests.

## Data and materials availability

Data and materials are provided upon reasonable request for the corresponding author.

## Notes

### Competing Interest Statement

The authors have declared no competing interest.

