## Supplemental infomation for "Systemic metabolic correlates of environmental sensitivity in group-housed mice"

**Table. S1 Detailed definition of behavioral metrics**

| Behavioral metrics of each individual | Definition |
|---|---|
| Activity state | -Defined using Gaussian mixture modeling (GMM; 4 components)<br>-Applied to z-scored features binned in 10-s intervals:<br>• Movement ratio<br>• Stop ratio<br>• Burst ratio<br>• Total distance |
| Spatial entropy | -Quantifies spatial dispersion of occupancy<br>-Computed as Shannon entropy across a 6 × 8 grid:<br>$-\sum p_i \log p_i$<br>($p_i$: occupancy probability of each grid cell) |
| Area occupancy | -Proportion of time an individual occupies its most frequently used grid cell<br>-Calculated relative to total observation time across all cells |
| Social distance during inactive state | Average pairwise distance between other individuals during inactive state |

| Dyadic interactions | Definition |
|---|---|
| Contact | -Pairwise distance ≤ 2 cells across consecutive frames<br>-Duration classified as *brief* or *sustained* using a 2-component GMM |
| Approach | -Distance to the partner decreases relative to the immediately preceding frame<br>-Evaluated at the onset of a contact event |
| Leave | -Distance to the partner increases relative to the immediately preceding frame<br>-Evaluated at the end of a contact event |
| Invade-evade | *Invade*<br>-Focal individual moves toward the target<br>$d_{AB}(t_{n-1}) > d_{AB}(t_n)$<br>-Distance moved by the focal individual > 2 cells<br><br>*Evade*<br>-Target individual moves away from the focal individual<br>$d_{BA}(t_{n-1}) < d_{BA}(t_n)$<br>-Distance moved by the target individual > 2 cells<br><br>*Invade–evade event*<br>-Invade and evade occur within ±1 frame<br>-One or more consecutive frames satisfying this condition are grouped<br>-Each grouped sequence is counted as a single event<br>-Events classified as *avoid* or *chase* are excluded |

| | |
|---|---|
| Chase | -Focal individual moves toward the target<br>$$d_{AB}(t_{n-1}) > d_{AB}(t_n)$$<br>-Distance moved by the focal individual > 1<br>-Cosine similarity between movement directions of focal and target > 0.7<br>-Pairwise distance < 3 cells<br>-Two or more consecutive frames satisfying these conditions are counted as a single event (Tsuchiya et al., unpublished) |
| Avoid | -Focal and target individuals are in proximity (pairwise distance < 2 cells, duration < 3 seconds)<br>-Distance moved by the focal individual > 5 cells<br>-Acceleration of the focal individual > 2 cells<br>-Target individuals are in a hyperactive state<br>-Target individuals do not move or do not approach the focal<br>-Pairwise distance increases up to 5 cells or more |
